# B-1 lymphoid cells develop independently of Notch signaling during mouse embryonic development

**DOI:** 10.1101/2020.11.09.375634

**Authors:** Nathalia Azevedo, Elisa Bertesago, Ismail Ismailoglu, Michael Kyba, Michihiro Kobayashi, Andrea Ditadi, Momoko Yoshimoto

## Abstract

The *in vitro* generation from pluripotent stem cells (PSCs) of different blood cell types, in particular those that are not replenished by hematopoietic stem cells (HSCs) like fetal-derived tissue-resident macrophages and innate-like lymphocytes, is of a particular interest. In order to succeed in this endeavor, a thorough understanding of the pathway interplay promoting lineage specification for the different blood cell types is needed. Notch signaling is essential for the HSC generation and their derivatives, but its requirement for tissue-resident immune cells is unknown. Using mouse embryonic stem cells (mESCs) to recapitulate murine embryonic development, we have studied the requirement for Notch signaling during the earliest B-lymphopoiesis and found that *Rbpj*-deficient mESCs are able to generate B-1 cells. Their Notch-independence was confirmed in *ex vivo* experiments using *Rbpj*-deficient embryos. In addition, we found that upregulation of Notch signaling was needed for the emergence of B-2 lymphoid cells. Taken together, these findings indicate that control of Notch signaling dosage is critical for the different B-cell lineage specification and provides pivotal information for their *in vitro* generation from PSCs for therapeutic applications.

## Introduction

It is well accepted that Notch signaling is indispensable for the generation of hematopoietic stem cell (HSC) from hemogenic endothelial cells (HECs) in the mouse embryo^1,2^, in particular, for the endothelial-to-hematopoietic transition (EHT) yielding HSCs. However, it remains unclarified the role of Notch signaling in HSC-independent hematopoietic programs, such as those generating tissue-resident immune cells (e.g. microglia, epidermal γδT-cells, and B-1a cells)^3^. Experiments in different animal models have shown that erythroid-myeloid progenitor (EMP) development in the yolk sac (YS) is largely unaffected in the absence of Notch signaling^1,2,4,5^, but little is known about embryonic lymphopoiesis. While T-cell lineage specification requires continuous Notch activation, B-cell development from adult HSCs occurs in the absence of active Notch signaling^6^. Whether the emergence of fetal B-1 cells is HSC-dependent is still controversial^7–9^ and therefore its requirement of Notch activation is currently unknown. Here we used *Rbpj*^*−/−*^ mouse embryonic stem cells (mESCs)^10^ and mouse embryos, defective for the canonical Notch signaling target, to determine the Notch signaling requirement in fetal B-lymphopoiesis. Our results indicate that not only Notch signaling is dispensable for the emergence of fetal B-1 cells, thus, indicating their HSC-independency, but also its fine tuning is critical for the emergence of B-2 cells.

## Methods

*Rbpj*^*+/−*^ and *Rbpj*^*−/−*^ mESCs were a kind gift from Dr. Timm Schroeder^10^. The primitive and definitive colony forming cells (CFCs) were produced from ESCs through embryoid body formation as previously reported^11,12^. T- and B-lymphoid cells were differentiated on OP9 or OP9-DL1 stromal cells with added 10ng/ml SCF and 10ng/ml IL7 as previously described^13^. VE-cadherin Cre mice^14^ were obtained from Dr. Nancy Speck. *Rbpj*-flox mice^6^ were obtained from Dr. Tasuku Honjo. The modified B-progenitor CFC assays were performed as previously reported^15^. AA4.1^+^CD19^+^ B-progenitors were harvested from the co-culture of mESC Flk-1^+^ with OP9 and were transplanted into the peritoneal cavity of sublethally (100rad) irradiated NSG neonates at day 1 or 2^16^.

## Results and Discussion

We first validated that *Rbpj*^*−/−*^ mESC hematopoietic differentiation faithfully phenocopies what has been reported in several animal models defective for Notch signaling^1,2,4^. Both day 3.25 and day 5.5 Flk-1^+^ cells from *Rbpj*^*−/−*^embryoid bodies (EBs) produced a higher number of classical defined primitive erythroid colony forming cells (EryP-CFCs) than that of *Rbpj*^*+/−*^ ESCs (Figure S1A-D, Figure S2A-D). In day 5.5 Flk-1^+^ primitive erythroid progenitor cells segregated to the CD41^+^ fraction that was more abundant in *Rbpj*^*−/−*^ than in *Rbpj*^*+/−*^ EBs (Figure S2E-G), confirming that primitive erythropoiesis is not properly terminated in the absence of *Rbpj*^*4,17*^.

We next tested the ability of day 5.5 Flk-1^+^ hemogenic endothelial cells (HECs) yielding EMP-like cells^12^ to undergo EHT, in serum free adherent culture conditions^12^ (Figure S3A) and found that *Rbpj*^*−/−*^ ECs generated less CD45^+^ cells and CFCs (2.9-fold) (Figure S3B-D). Gene expression analysis via qPCR revealed that known Notch targets^2^ (*Hes1* and *Gata2*) as well as pivotal hematopoietic transcription factors (*Tal1* and *Runx1c*) are significantly down-regulated in day 5.5 Flk-1^+^ HECs (Figure S3E). These results, in line with those obtained with chimeric *Notch1^−/−^* mice^4^, suggest that *Rbpj* is not required for the generation of EMP hematopoiesis, but alters the proliferation of hematopoietic progenitors. Alternatively, day 5.5 Flk-1^+^ cells, and probably EMP HECs in the mouse embryos, are heterogenous and comprise both Notch-dependent and -independent precursors. Collectively, these data show that *Rbpj*^*−/−*^ mESCs can be used to dissect the Notch signaling requirement of different hematopoietic embryonic progenitors.

We then investigated the lymphoid potential of *Rbpj*^*−/−*^ ESCs. In OP9-DL1 co-culture,^13^ while *Rbpj*^*+/−*^ Flk-1^+^ cells differentiated into CD4^+^CD8^+^ double positive (DP) T-cells, *Rbpj*^*−/−*^ Flk-1^+^ showed maturation arrest at DN1 stage and lineage switch into CD19^+^ B-cells (Figure S3F-G) consistent with the previous report of conditional *Rbpj* knockout mice^6^. This is also compatible with a recent report showing that the first embryonic thymopoiesis-initiating progenitors are present in the absence of *Rbpj*^18^, but are unable to progress beyond the DN1, although this report did not assess the alternative B-cell fate. The earliest thymic T-progenitors (ETPs) in DN1 have been reported to possess B- and myeloid potential^19^ and it is well accepted that lineage choice of T- or B-cells depends on Notch signaling^6,20^. Therefore, our observation of alternative *Rbpj*^*−/−*^ B-cell specification in T-cell cultures supports a model where ETP-like cells are produced and switch to B-cell fate in the absence of Notch signaling.

Next, we confirmed B-cell production from *Rbpj*^*−/−*^ Flk-1^+^ cells in the OP9 co-culture, at a similar level to heterozygous ESCs. (Fig. 1A, B). We recently reported that mESC-derived B-cells engrafted immunodeficient neonates as peritoneal B-1 cells^16^. Likewise, we injected AA4.1^+^CD19^+^ B-cells differentiated from *Rbpj*^*+/−*^ and *Rbpj*^*−/−*^ ESCs into NSG neonates. Indeed, *Rbpj*^*−/−*^ B-1 cells, but not B-2 cells, were engrafted in the recipient peritoneal cavity (Fig. 1C, D). Thus, *Rbpj*^*−/−*^ ESCs can differentiate into transplantable B-1 cells.

**Fig. 1.**
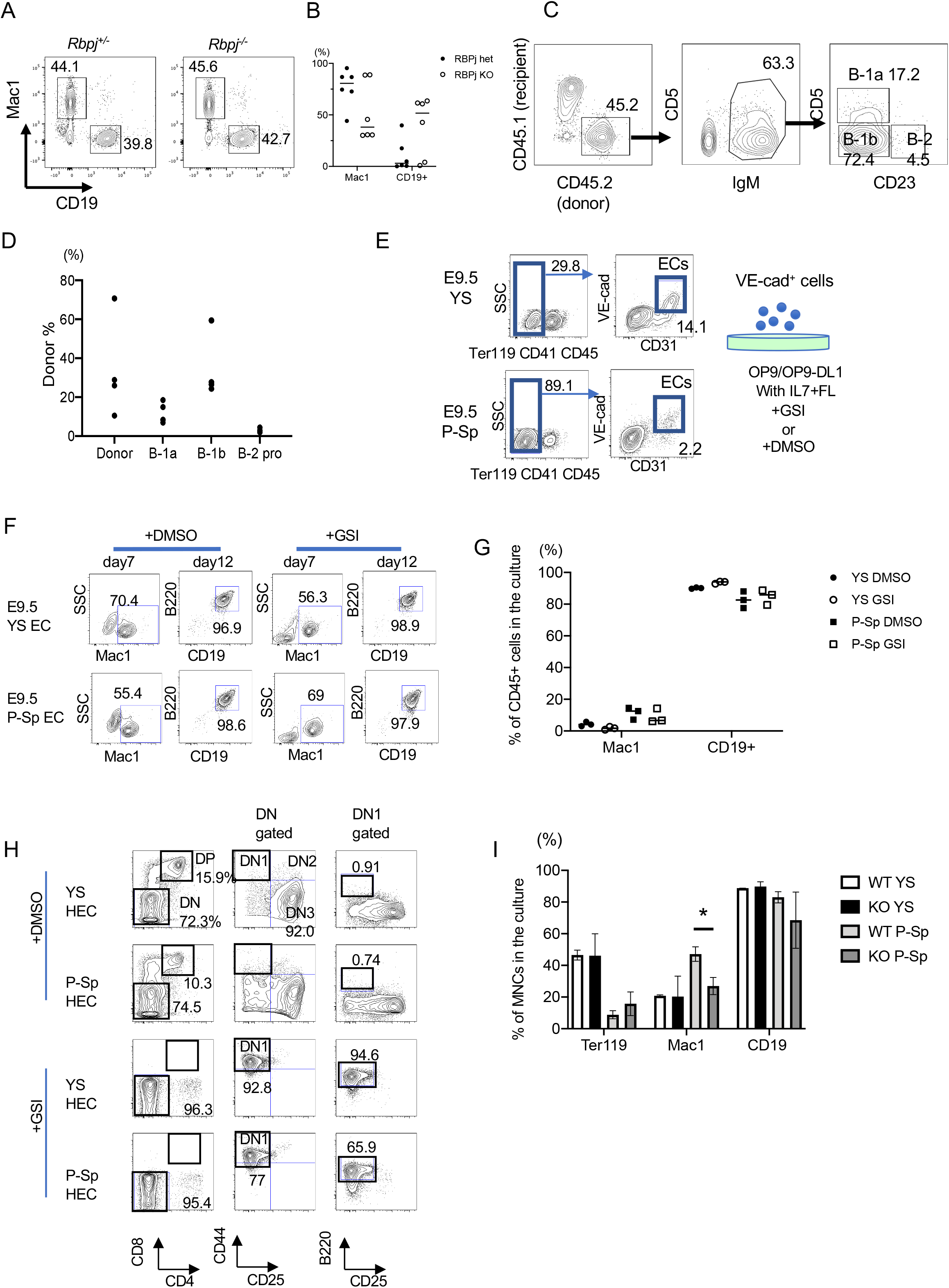
*Rbpj*^*−/−*^ ESCs can differentiate into erythro-myeloid, B-cells, and early lymphoid progenitors in vitro. A, Representative FACS plots of Mac1^+^ myeloid and CD19^+^ B-cells produced from Flk-1^+^ cells on OP9 co-culture are shown. B, The percentages of Mac1^+^ and CD19^+^ cells among CD45^+^ cells in the ESC culture with OP9 are shown (n=6). C, Representative FACS plots and the donor derived cells of the peritoneal cells of the recipient NSG mice transplanted with *Rbpj*^*−/−*^ ESC-derived B-1 cells are depicted (n=4). E, Sorting strategy of CD31^+^VE-cadherin^+^ HECs from E9.5 YS and P-Sp is shown. More than three times experiments were performed. F and G, Representative FACS plots and percentages among CD45^+^ cells of the supernatant of HEC co-culture with OP9 with/without GSI are depicted (n=3). H, The representative FACS plots of YS/PSP co-culture with OP9-DL1 with/without GSI are depicted (n=3 for each group). I, The percentages of each blood lineages among mononuclear cells (MNCs) in the supernatant of WT and *VCCre:RbpjKO* YS and P-Sp co-culture with OP9 are shown (n=3 for each group). The percentage of Ter119^+^ and Mac1^+^ cells was examined at day 6 and the percentage of B-cells was examined at day10 of OP9 co-culture. Student’s unpaired t-test *p < 0.05.

Next, we tested the effect of Notch signal inhibition on HECs of the mouse embryo. FACS sorted VE-cadherin^+^ (VC^+^) ECs from E9.5 YS and paraaortic splanchnopleura (P-Sp) were plated on OP9 cells with or without gamma secretase inhibitor (GSI) that inhibits the cleavage of Notch intracellular domain (NICD), thus inhibits Notch-intracellular signaling^21^ (Fig. 1E). Mac1^+^ myeloid and CD19^+^B220^+^ B-cells were similarly produced regardless of GSI addition (Fig. 1F, G). In OP9-DL1 culture with GSI, the T-cell development from VC^+^ ECs was arrested at DN1 stage, where B220^+^ B-cells were detected, whereas DN2 or DN3 committed T-cells were produced in control culture (Fig. 1H). We also confirmed the Notch-independent B-cell potential using *VCCre:Rbpjf/f* embryos (*Cre^+^RbpjKO*). The YS and P-Sp cells from *VCCre*^*+*^*RbpjKO* and control embryos displayed similar erythro-myeloid and B-cell production (Fig. 1I). These results indicate the presence of Notch signal independent B-lymphopoiesis.

Next, the positive effect of Notch signaling on the HECs was examined using Doxycycline (Dox) inducible NICD-overexpressing (iNICD) ESCs^22^. iNICD Flk-1^+^ cells were differentiated on OP9 with 0, 100, and 500 ng/ml Dox. Flk-1^+^ cells produced CD19^+^ B-cells without Dox and CD4^+^CD8^+^ DP and DN CD25^+^ T-committed cells with 500ng/ml Dox (Fig. 2A). Interestingly, iNICD Flk-1^+^ cells produced both T- and B-cells in the same well with 100ng/ml Dox. We hypothesized that the HEC fate into B-1, B-2, and T-cell lineage is determined by the dosage of Notch signaling as it is also seen in B- and T-cell differentiation from BM HSPCs^20^. Additionally, we and otheres showed that Notch signaling induced B-2 potential in HSC-precursors^9,15,23^. The modified B-progenitor CFC assays showed more B-1+B-2 progenitor colonies with a moderate dosage of Dox (Fig. 2B, C). These results indicate that fine tuning of Notch-signaling control B-1 and B-2 cell fate specification from the HECs.

**Fig. 2.**
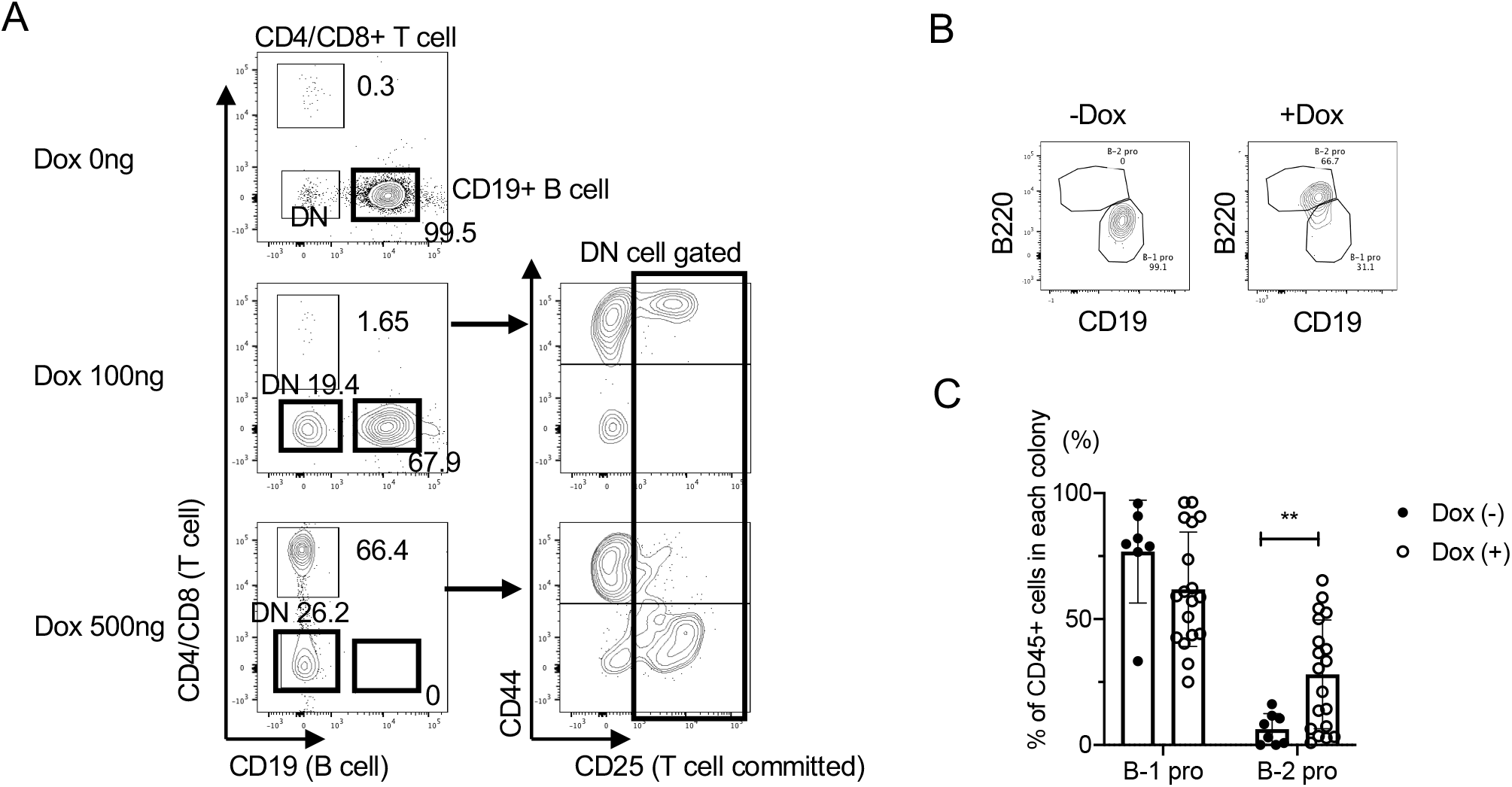
The different dosages of Notch signaling affect the lineage choice of HECs between B-1 and B-2 cells. The representative FACS plots of Flk1+ cell-coculture with OP9 with different dosages of doxycycline (n>3). B, The representative FACS plots of B-1 and B-2 progenitors that form a single colony is depicted. C, B-1 and B-2 progenitor percentages among CD45^+^ cells in each colony with and without doxycycline are shown. Dox (−) n=8, dox (+) n=19. Student’s unpaired t-test **p < 0.01.

Here, using *Rbpj*^*−/−*^ ESC and mouse models, we demonstrated that the development of the earliest lymphoid progenitors, in particular of B-1 cells, can occur in the absence of Notch signaling and therefore independently of HSCs. Because B-1a cells are mainly fetal derived and are not replenished by BM progenitors, the conditioning regimen for BM transplantation therapy may induce permanent B-1 cell deficiency in humans, which may cause patients' susceptibility to certain bacterial/viral infections^24^. Additionally, natural IgM antibodies produced by B-1 cells have protective roles against atherosclerosis and other inflammatory diseases^25^. Therefore, there is an unmet clinical need for B-1 re-establishing cell therapies. Our findings are critical for the future development of strategies for the *in vitro* generation of B-1a cells from PSCs.

## Acknowledgment

This study is supported by NIAID R01AI121197 (M.Y.) and Fondazione Telethon, Italy (SR-TIGET Core Grant, Project C4) (A.D.).

## Authorship and conflict of interest

N.A., E.B., and M.K. (UTHealth) performed the experiments, I.I. and M.K. produced and provided the reagents, A.D. and M.Y. conceived the study, performed the experiments, analyzed the data, and wrote the manuscript. All authors declear no competing financial interests.

---

**Fig. S1.**
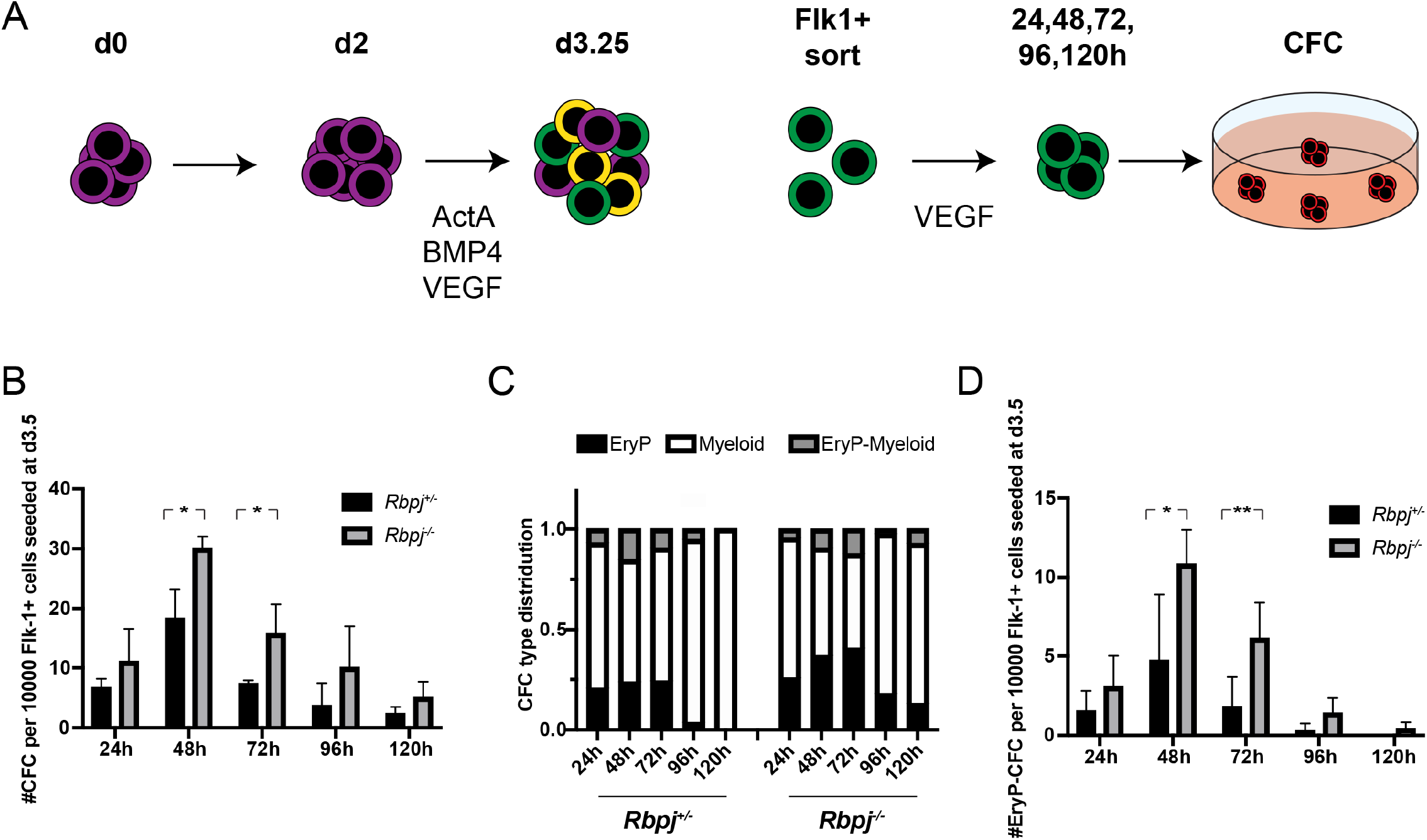
Primitive erythropoiesis termination is impaired in the absence of Notch signaling. A, Schematic representation of the mESC differentiation for the generation of the primitive hematopoietic program. B, Kinetics of CFC development from Flk-1^+^-derived aggregates and their hematopoietic lineage distribution (C). D, Kinetics of EryP-CFC. n=3, independent experiments. Student’s unpaired t-test *p < 0.05, **p < 0.01.

**Fig. S2.**
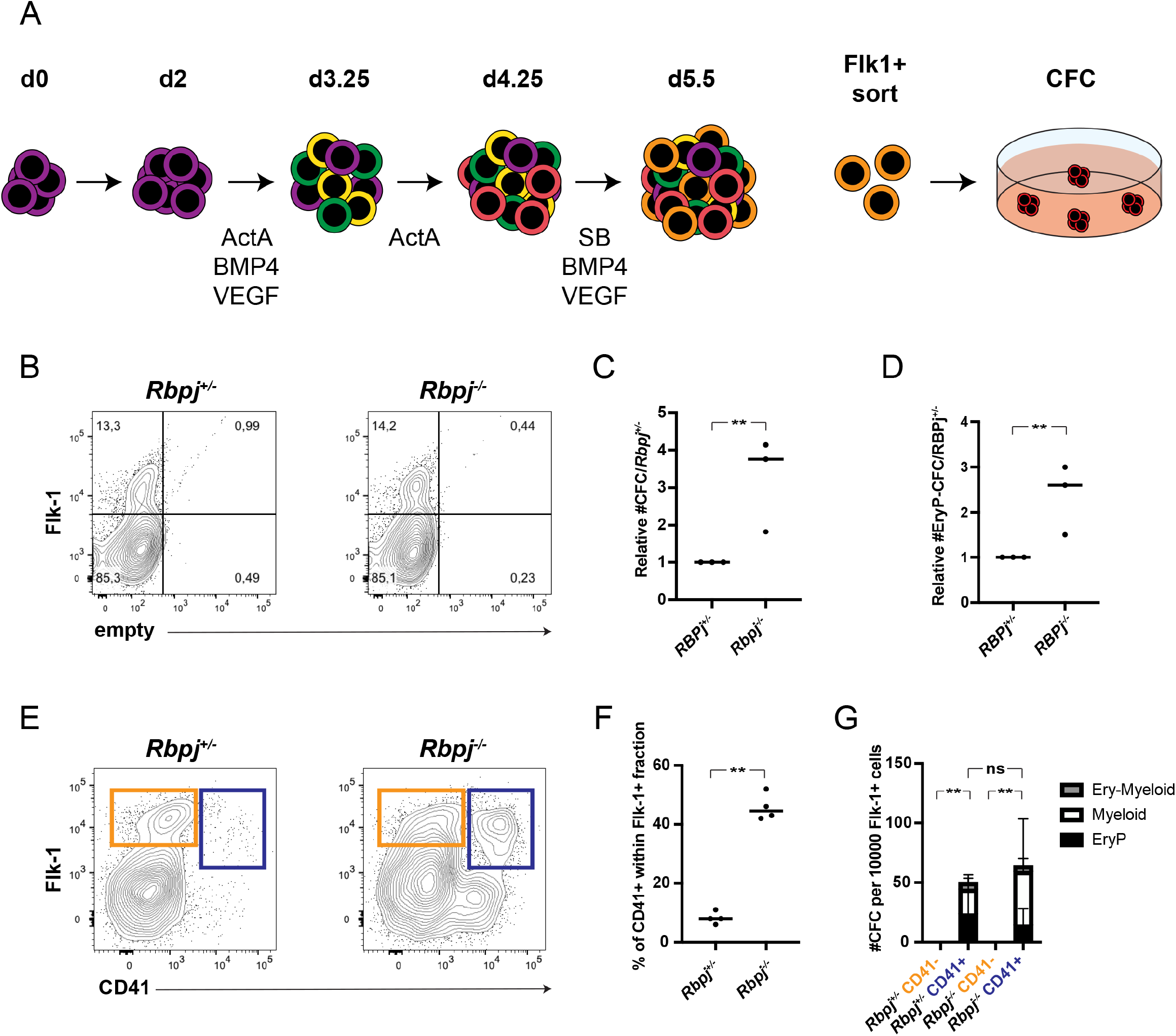
Residual primitive erythropoietic progenitor segregate to CD41^+^ cells in d5.5 differentiating cultures. A, Schematic representation of the mESC differentiation for the generation of the EMP like hematopoietic program. B, Representative flow cytometric analysis of the Flk-1 expression in d5.5. EBs, of 3 independent experiments. C, Relative number of CFCs and (D) EryP-CFC generated from d5.5 Fk-1^+^ cells. E, Representative flow cytometric analysis of the Flk-1 and CD41 expression in d5.5. EBs. F, Quantification of CD41^+^ cells present within the Flk1^+^ population. G, Frequency of CFC within the CD41 fractions of Flk-1^+^ cells. n=3, independent experiments. Student’s unpaired t-test *p < 0.05, **p < 0.01.

**Fig. S3.**
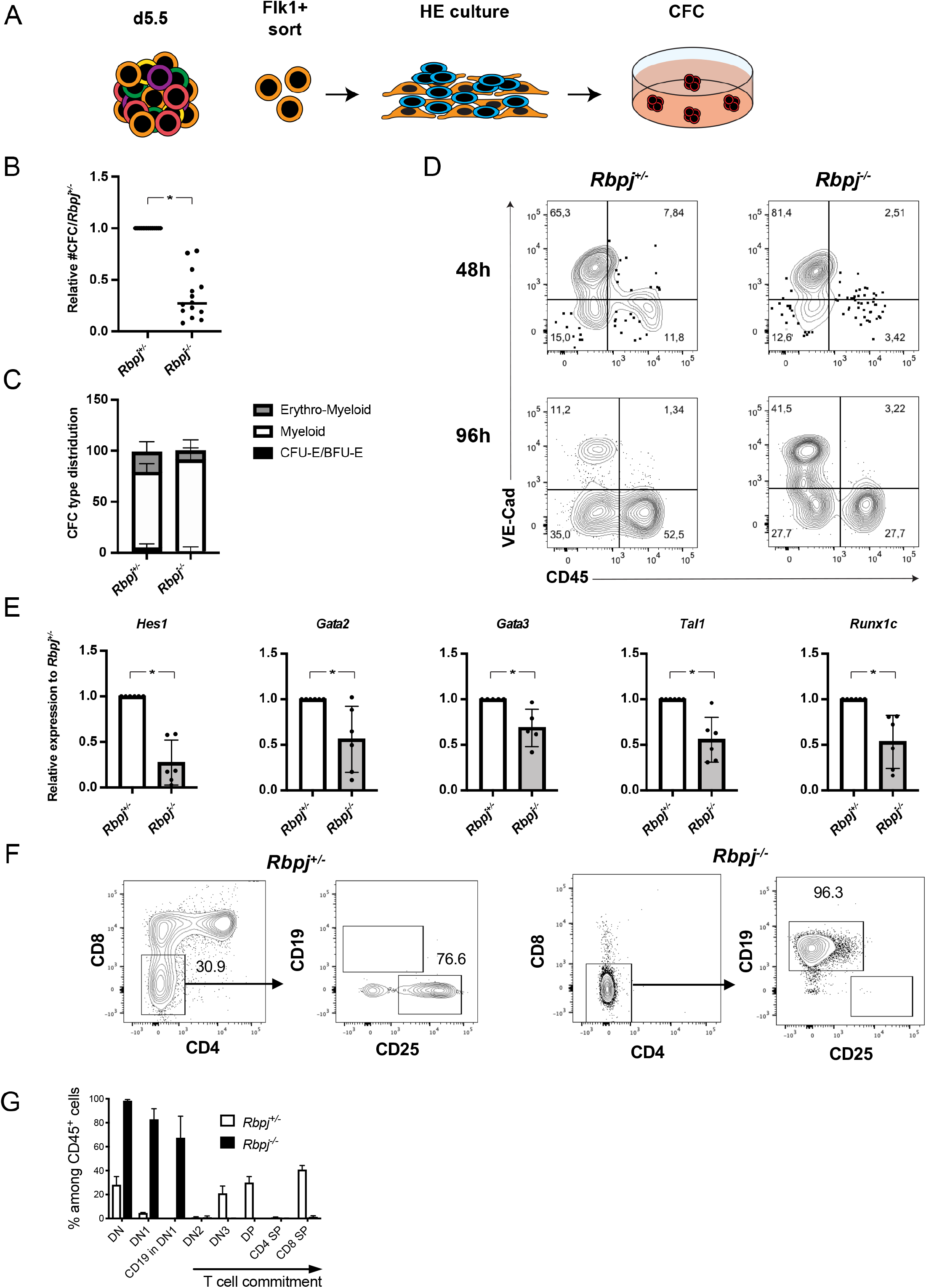
Hematopoietic output from HECs yielding EMP-like progenitors is reduced but not abrogated in the absence of Notch signaling. A, Schematic representation of the mESC differentiation for the HEC cultures. B, Representative flow cytometric analysis of the VE-Cad and CD45 expression after 48 and 96 hours of HEC culture. C, Relative number of CFCs obtained after 96 hours of HEC culture and (D) their lineage distribution. E, qRT-PCR-based gene expression analysis in d5.5 Flk-1^+^ cells. F, Representative flow cytometric analysis of lymphoid makers in cells obtained from *Rbpj*^*+/−*^ and *Rbpj*^*−/−*^ Flk-1^+^ cells cultured on OP9-DL1 for 15 days. G, Quantification of the proportion of each T-cell stage of Flk-1^+^ cell differentiation on OP9-DL. DN (Double negative): CD4^−^CD8^−^, DN1: CD4^−^CD8^−^CD25^−^CD44^+^, DN2: CD4^−^CD8^−^CD25^+^CD44^+^, DN3: CD4^−^CD8^−^CD25^+^CD44^−^, SP: single positive. N>3, independent experiments. Student’s unpaired t-test *p < 0.05, **p < 0.01.

